# Meta-analysis identifies microbial signatures of disease in murine models of inflammatory bowel disease

**DOI:** 10.1101/515205

**Authors:** Sudipta Saha, Alberto Martin, William Wiley Navarre

**Affiliations:** Department of Global Health and Population, Harvard T. H. Chan School of Public Health, Boston, MA, USA; Department of Immunology, University of Toronto, Toronto, ON, Canada; Department of Molecular Genetics, University of Toronto, Toronto, ON, Canada

## Abstract

The gut microbiota plays a central role in modulating intestinal inflammation, but the identification of specific inflammation-associated microbes has remained elusive. Here, we perform a meta-analysis on metagenomic data from 12 different studies of murine colitis triggered by a variety of genetic and environmental factors with the goal of finding bacterial taxonomic groups that can act as signatures of health or disease across studies, and that can be used to discriminate between healthy and diseased mice. We leveraged recent developments in 16S analysis tools to identify amplicon sequence variants (ASVs) instead of the traditional Operational Taxonomic Units, and used the EZTaxon reference database that distinguishes between currently unnamed and uncharacterized 16S phylotypes. Random Forest model and differential abundance analysis were used to detect microbial signatures that could consistently differentiate healthy from diseased mice, and a ‘dysbiosis index’ was constructed from these. This dysbiosis index was able to correctly distinguish samples derived from inflamed and non-inflamed mice in the majority of studies and significantly outperformed other frequently used metrics of dysbiosis including alpha-diversity, proteobacterial abundance, and the ratio of Bacteroidetes to Firmicutes. 10 of 12 bacteria we identify as associated with the diseased state are members of the order Bacteroidales, including several species from the abundant but poorly understood S24-7 family. The implications of these findings are discussed.

## INTRODUCTION

The human gut contains vast numbers of bacteria, viruses and fungi that collectively make up the gut microbiota, which plays a pivotal role in the host’s health (1). While the microbiome is important for fundamental host processes like digestion (2), metabolism (3) and immune system development (4), changes in the microbiome, often termed dysbioses (5), have been linked to several diseases, including inflammatory bowel disease (IBD) (6). IBD is a group of relapsing and remitting inflammatory disorders that mainly manifest as Crohn’s disease or ulcerative colitis (7, 8). Human studies have mostly been retrospective cohort ones, which limits their utility in elucidating causal links between changes in the microbiome and disease onset. Regardless, understanding these changes remain important, especially if there is a particular “dysbiotic” microbiome, or microbiome members, associated with disease. Although 16S sequencing of the bacterial microbiota have allowed comprehensive investigations of such changes (9), inter-individual variability within studies, and a lack of standardized techniques across studies (differing extraction and sequencing protocols, 16S variable regions, analysis pipelines and taxonomic reference databases) hinders comparisons of large sample sets to find consistent microbial signatures of disease (10). In order to compare and synthesize results from different human studies, meta-analyses, starting from the original sequencing data, have been conducted, resulting in the discovery of consistent microbial signatures in IBD patients (11, 12). Such meta-analyses have significant potential for designing non-invasive sequencing based diagnostic tools for IBD onset (12). They also reveal interesting insights into the relationship between the microbiota and disease; for example, meta-analyses of microbiome studies investigating obesity and IBD suggest that although metrics like the Bacteroidetes to Firmicutes ratio and alpha diversity may be significant in some studies, they do not seem to be consistent across studies (11, 13). In spite of the contributions of human meta-analyses to clinical diagnosis and broad inferences, causal and mechanistic inference remains challenging.

To shed mechanistic insight on the link between IBD and the microbiota, murine models have been relied upon heavily (6, 14, 15). Yet, despite the professed advantages of reproducibility and well-defined conditions, murine samples also seem to have significant inter-individual variability, lack of standardization across studies, and sample sizes are often small (14, 16). Furthermore, there are multiple widely-used mouse models for IBD, including Dextran Sodium Sulfate (DSS) induced colitis, 2,4,6-trinitrobenzenesulfonic acid (TNBS) induced colitis, T-cell adoptive transfer, and Interleukin-10 deficiency (17, 18). Studies investigating the role of various immunomodulatory genes in IBD also often report changes in the microbiota (19, 20).

The existing literature mostly consists of descriptions of microbiota changes associated with colitis in various mice models (19, 21, 22), with only some assessing colitogenic potential of particular microbiotas, or honing in on particular microbes (23, 24). The lack of standardization prevents meaningful comparison of the changes reported, and the question remains as to whether there are any consistent markers of inflammation in mice models. Reviews published have been descriptive (25, 26). Since there is little overlap between the human and murine microbiota, host-specific analysis is paramount (27). In the context of a poorly catalogued murine microbiota with limited cultured isolates (28), identification of a microbial signature can help focus isolation efforts and mechanistic studies on the best microbial candidates for further research in mouse models.

Here we report a meta-analysis of 12 studies/datasets that utilize 16S sequencing to describe a link between development of colitis and changes in the microbiota in murine models. We aimed to find bacterial taxonomic groups that are consistent signatures of health and disease across studies, and that can be used to discriminate between healthy and diseased mice. We only included studies whose raw 16S sequencing read files were available and thus were able to standardize the analysis and directly compare the studies. We leveraged recent developments in 16S analysis tools to identify amplicon sequence variants (ASVs) instead of the traditional Operational Taxonomic Units (OTUs) (29), and used the EZTaxon reference database that distinguishes between currently unnamed and uncharacterized 16S phylotypes (30). We used a Random Forest model and differential abundance analysis to detect any consistent microbial signatures that differentiate healthy from diseased mice, and constructed a “dysbiosis index” from these. Finally, since alpha diversity, Bacteroidetes-to-Firmicutes (BF) ratio and Proteobacteria levels are often used as markers of microbiome health, we investigated the utility of these in discriminating colitic from healthy samples (11, 13, 31).

## RESULTS

### Study Search, Inclusion and Data Aggregation

To identify studies that investigated the murine intestinal microbiota in the context of intestinal inflammation, we conducted a systematic search of NCBI PubMed for articles that contained terms relating to microbiota, intestinal inflammation and murine models in the title and abstract, published between 2012 and 2016, and was not a review. We followed the preferred reporting items for systematic reviews and meta-analyses (PRISMA) guidelines to limit inclusion bias (32). We screened the title and abstracts of 816 articles yielded by the search for eligibility, and obtained 2 additional studies from knowledge of published literature and our own currently unpublished dataset (Martin A. et al. unpublished) (819 total); 79 full-text articles were then assessed; 44 were retained to be checked for data availability; 10 had publicly available data and 2 provided access by the time of publication (19–22, 33–39) (**Table 1, Fig 1**).

**FIG 1:**
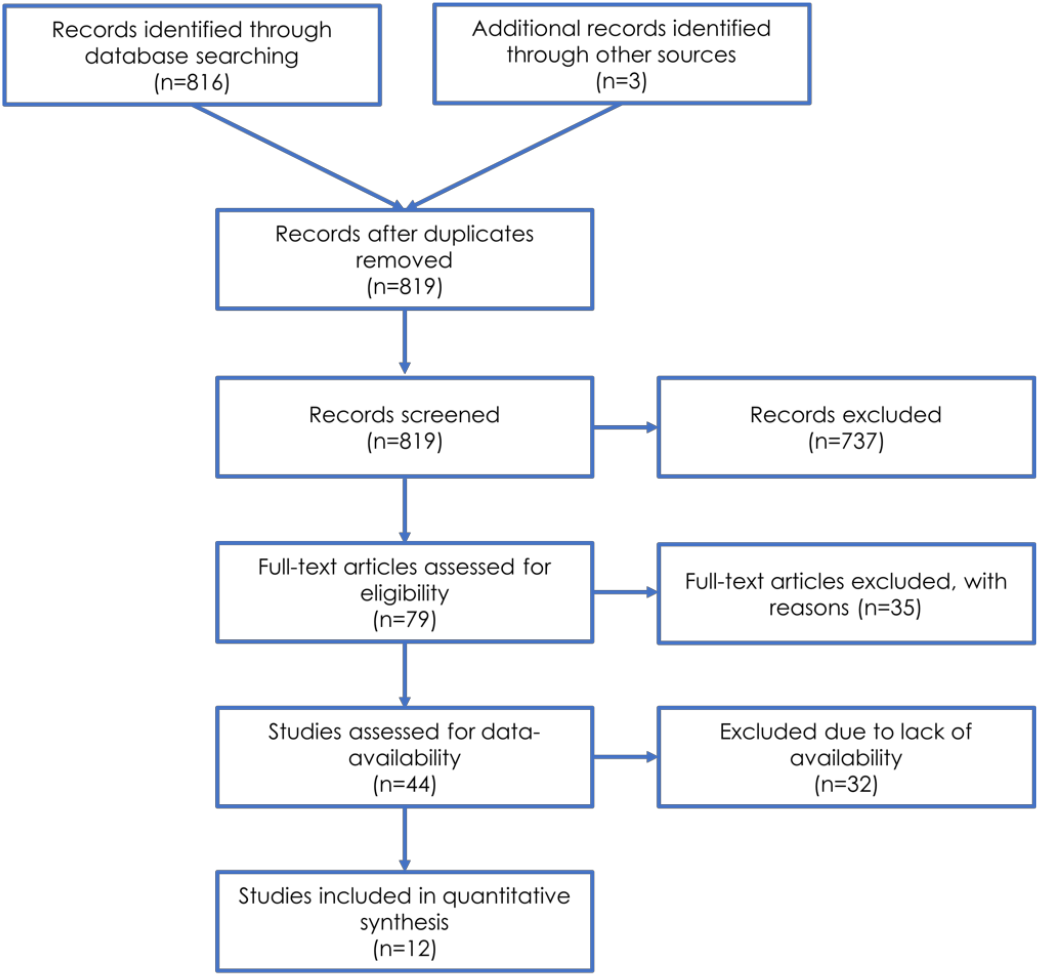
Flow diagram of literature search and review for inclusion in meta-analysis, represented according to PRISMA guidelines.

**TABLE 1.** Summary of studies included in the meta-analysis. DSS = Dextran Sulfate Sodium, IL10 = Interleukin 10, NSAID = Non-Steroidal Anti-Inflammatory Drug, TNBS = 2,4,6-trinitrobenzene sulfonic acid.

| Study, Year | Title | Colitis Model | Mouse Strains | DNA extraction | Region | Platform | Biological Source | Sample counts | Data Source |
| --- | --- | --- | --- | --- | --- | --- | --- | --- | --- |
| Jacobs et al. 2017<sup>19</sup> | Microbial, metabolomic, and immunologic dynamics in a relapsing genetic mouse model of colitis induced by T-synthase deficiency. | Tsyn mice (C1galt1 or T-synthase deleted) | C57BL/6 | MO Bio PowerSoil DNA Isolation Kit | V4 | Illumina HiSeq | Intestinal lavage and wash | 54 Healthy<br>33 Colitis | Public, PRJNA318692 |
| Chassaing et al. 2015<sup>20</sup> | Dietary emulsifiers impact the mouse gut microbiota promoting colitis and metabolic syndrome. | Dietary emulsifiers in IL10-/- mice | C57BL/6 | MO Bio PowerSoil DNA Isolation Kit | V4 | Illumina MiSeq | Fecal Samples | 175 Healthy<br>9 Colitis | Public, PRJEB8035 |
| Laubitz et al. 2016<sup>21</sup> | Reduced Epithelial Na<sup>+</sup>/H<sup>+</sup> Exchange Drives Gut Microbial Dysbiosis and Promotes Inflammatory Response in T Cell-Mediated Murine Colitis. | NHE3-/-, Rag2-/- | 129S6/SvEv | Beads + Proteinase, Phenol-chloroform | V4 | Illumina MiSeq | Fecal Samples | 76 Healthy<br>16 Colitis | Public, 10.17605/OSF.IO/UWFAP |
| Yeom et al. 2016<sup>22</sup> | Sasa quelpaertensis leaf extract regulates microbial dysbiosis by modulating the composition and diversity of the microbiota in dextran sulfate sodium-induced colitis mice. | DSS | C57BL/6 | Fast DNA SPIN Kits | V1-V3 | Roche 454 | Fecal samples | 6 Healthy<br>6 Colitis | Public, PRJEB13815 |
| Lamas et al. 2016<sup>32</sup> | CARD9 impacts colitis by altering gut microbiota metabolism of tryptophan into aryl hydrocarbon receptor ligands. | DSS in CARD9-/- mice | C57BL/6J | Beads in FastPrep, isopropanol. | V3-V4 | Illumina MiSeq | Fecal samples | 6 Healthy<br>9 Colitis | Public, PRJEB9079 |
| Whitfield-Cargile et al. 2016<sup>33</sup> | The microbiota-derived metabolite indole decreases mucosal inflammation and injury in a murine model of NSAID enteropathy. | NSAID (indomethacin) | C57BL/6J | MO Bio PowerSoil DNA Isolation Kit | V4 | Illumina MiSeq | Fecal samples | 33 Healthy<br>3 Colitis | Public, PRJNA290483 |
| Martin, unpublished | Experiments showing IL10-/- deletion caused spontaneous colitis in some mouse cages but not others | DSS in IL10-/- mice | C57BL/6 | DNA from soil kit (Macherey-Nagel) | V4 | Illumina MiSeq | Fecal samples | 11 Healthy<br>17 Colitis | Request |
| Sassone-Corsi et al. 2016<sup>34</sup> | Microcins mediate competition among Enterobacteriaceae in the inflamed gut. | DSS | C57BL/6 Slc11a1<sup>+</sup> | QIAamp DNA Stool Kit | V4 | Illumina MiSeq | Fecal samples | 5 Healthy<br>7 Colitis | Public, PRJEB15700 |
| Moschen et al. 2016<sup>35</sup> | Lipocalin 2 protects from inflammation and tumorigenesis associated with gut microbiota alterations. | Lcn2-/-, IL10-/- | C57BL/6J | FastDNA SPIN Kit, Precellys 24 homogenizer | V1-V2 | Illumina MiSeq | Cecal content | 11 Healthy<br>19 Colitis | Public, ERP014639 |
| Berry et al. 2015<sup>36</sup> | Intestinal microbiota signatures associated with mice experiencing recurring colitis. | DSS | C57BL/6 | Phenol-chloroform, bead | V6-V9 | Roche 454 | Intestinal flush, fecal samples | 36 Healthy<br>36 Colitis | Request |
| Vereecke et al. 2014<sup>37</sup> | A20 controls intestinal homeostasis through cell-specific activities. | A20 deletion | C57BL/6 | QIAamp DNA Stool Mini Kit | V3-V5 | Roche 454 | Cecal content | 16 Healthy<br>6 Colitis | Request |
| He et al. 2016<sup>38</sup> | Dysbiosis of the fecal microbiota in the TNBS-induced Crohn's disease mouse model | TNBS | BALB/c | Phenol-chloroform, bead | V5-V4 | Ion Torrent PGM | Fecal samples | 5 Healthy<br>6 Colitis | Public, ERP011541 |

Search terms and screening and inclusion strategy are outlined in Methods. Briefly, we looked for studies that did a 16S sequence-based analysis of non-synthetic murine microbiota in various IBD models before and after onset of colitis. We excluded infection-based inflammation and samples administered antibiotics as these are likely to have changes that go well beyond colitis associated ones. Studies that only assessed microbiota before inflammation onset were excluded because we were interested in a microbial signature associated with the *onset* of colitis, rather than a microbiota that is associated with increased *susceptibility* to colitis.

Selection of relevant samples within the 12 studies yielded 601 samples, of which 434 were healthy and 167 had colitis. We used a standardized custom data-processing pipeline to detect Amplicon Sequence Variants (ASVs) using the DADA2 algorithm that leverages quality information from sequence reads for sequence inference. Taxonomy was assigned using a custom script and the EZTaxon database to be able to distinguish and name currently uncultured, but sequenced, phylotypes. Only classified taxons were kept and after filtering for rare taxa and merging datasets, we obtained 1558 unique taxonomic groups (detailed methods in Materials and Methods).

### Beta Diversity

Beta diversity, the between-sample diversity, can provide insight as to whether mice with colitis have a different microbial community structure compared to healthy mice (40). We calculated the Bray-Curtis distances (41) between samples and used Principal Coordinates Analysis (PCoA) to visualize the microbial communities. Plotting the samples on the first three coordinates suggested that samples tended to cluster by study more than by disease status (**Fig 2a, b**). However, a Permutational Analysis of Variance (PERMANOVA) (42) suggested that microbiome composition differed by both disease status and study (p < 0.001). Furthermore, within each study, PERMANOVA revealed significant differences due to disease status in all except one study (**Fig S1**). Visually, within each study, disease status often provided a stark differentiation in PCoA plots, which was not replicated in the pooled data. However, the significant PERMANOVA result suggested the possibility of differences between communities based on disease status, across studies.

**FIG 2:**
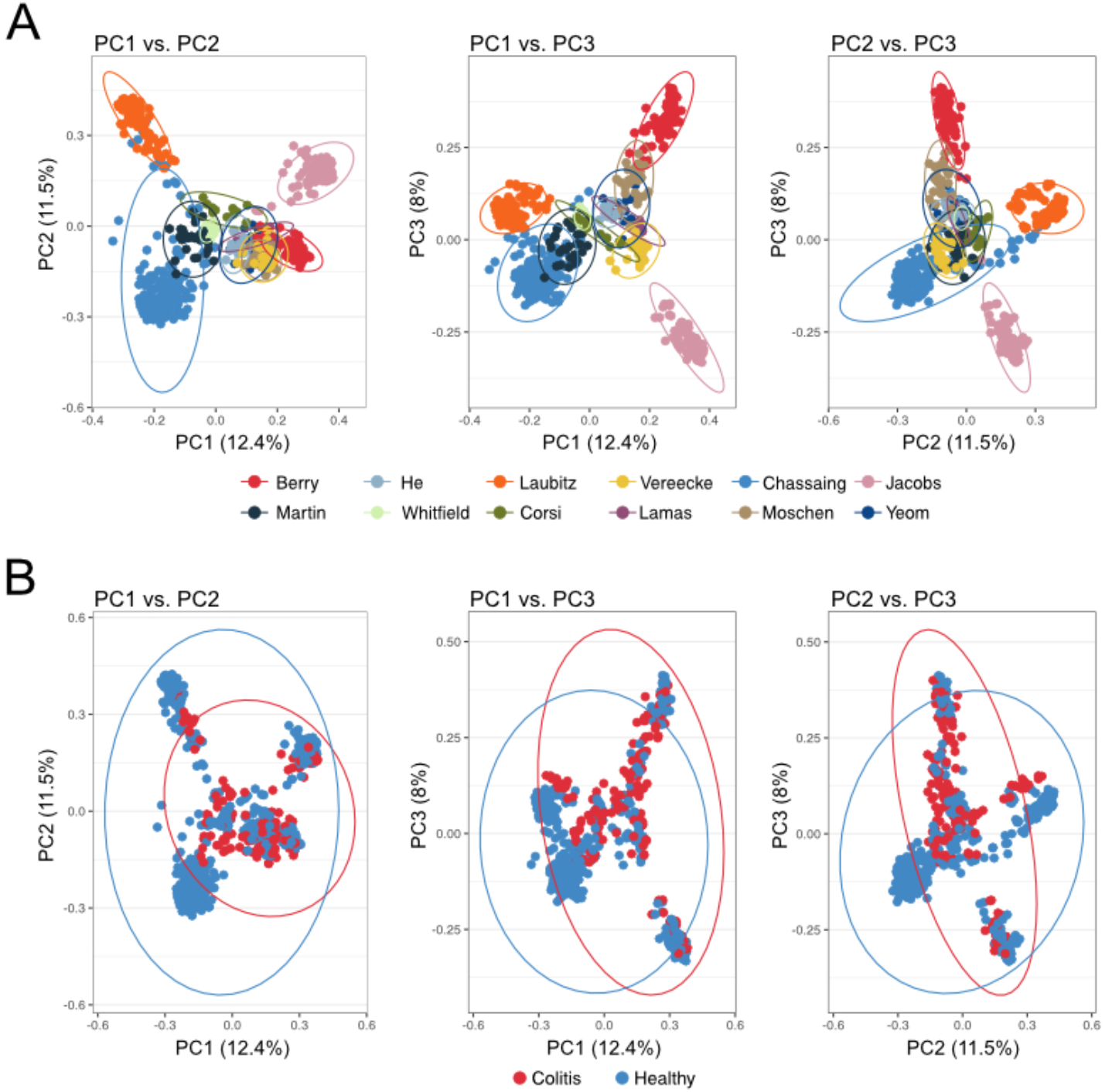
Principle Coordinates Analysis (PCoA) plots depicting the relationships between sample microbial compositions along the first three principal coordinates. Colored points and the associated ellipses distinguish between the different studies/datasets (A) or disease status (B).

### Alpha Diversity

Alpha diversity, the diversity of the microbiome within each sample, is regularly investigated as a marker of “health” in both human and mouse studies. While individual studies have found associations between reduced alpha diversity and obesity and IBD in humans, meta-analyses have found the evidence for such relationships to be weak (13, 43). Another oft-used, yet inconsistent, within-sample marker is the ratio of Bacteroidetes to Firmicutes ratio. Lastly, a bloom of Proteobacteria has also been associated with dysbiosis. We investigated the utility of using these markers in discriminating between healthy and diseased mice.

In our meta-analysis, we do not find a consistent relationship between alpha diversity and colitis in mice. Seven of the 11 studies had significant differences in the Shannon index (*H*) between healthy and diseased mice, with 5 having higher values of *H* (lower diversity) in healthy, and 2 having higher diversity in healthy (**Fig 3a**). Similarly, there was no consistent relationship between colitis and the Bacteroidetes to Firmicutes ratio; 2 studies had a significantly lower ratio in healthy, while 2 had a significantly higher ratio in healthy (**Fig 3b**). Finally, there was also no consistent relationships between disease status and the relative abundance of Proteobacteria (**Fig 3c**). For all three measures, the pooled data did not show a significant difference between the healthy and diseased mice when tested using random effects models (p > 0.05).

**FIG 3:**
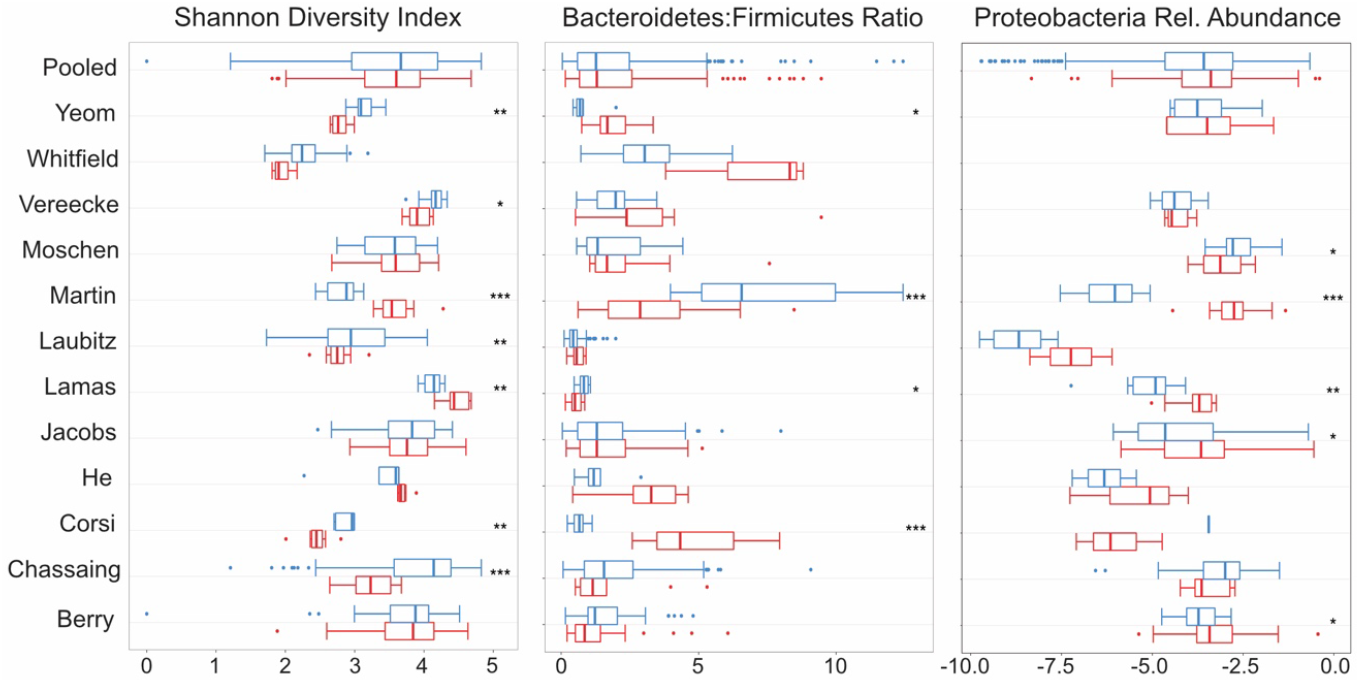
Boxplots of commonly used diversity and composition metrics. From left, each panel shows distribution and statistical significance of Shannon Diversity Index, Bacteroidetes-Firmicutes Ratio and Relative Abundance of Proteobacteria, within each study and for pooled samples. Blue represents healthy samples and red represents colitic samples. * = p < 0.05, ** = p < 0.01, *** = p < 0.001.

### Random Forest Models

Given the significant PERMANOVA results, inconsistent alpha diversity metrics, and the success of statistical learning techniques in previous studies, we hypothesized that a random forest model would be able to discriminate between diseased and healthy mice (12, 43, 44). A random forest model works by building hundreds of decision trees, with a cut-off value (abundance) for a particular feature (taxon) being selected at each split to maximize the correct classification of the outcome (disease status) among the samples being used for training. For a new sample, the trees are used to classify it as diseased or healthy (45). Model performance in discrimination can be summarized using the area under the receiver operating curve (AUROC), with 0.5 being as good as random, and 1 being perfect prediction. We built a cross-validated random forest model on a randomly selected 70% of the samples, which yielded an AUROC of 0.975. When this model was used to predict the remaining 30% of the samples, the AUROC was 0.972, suggesting a lack of overfitting and the presence of a detectable microbial signature of disease across studies included in this analysis (**Fig 4a**).

**FIG 4:**
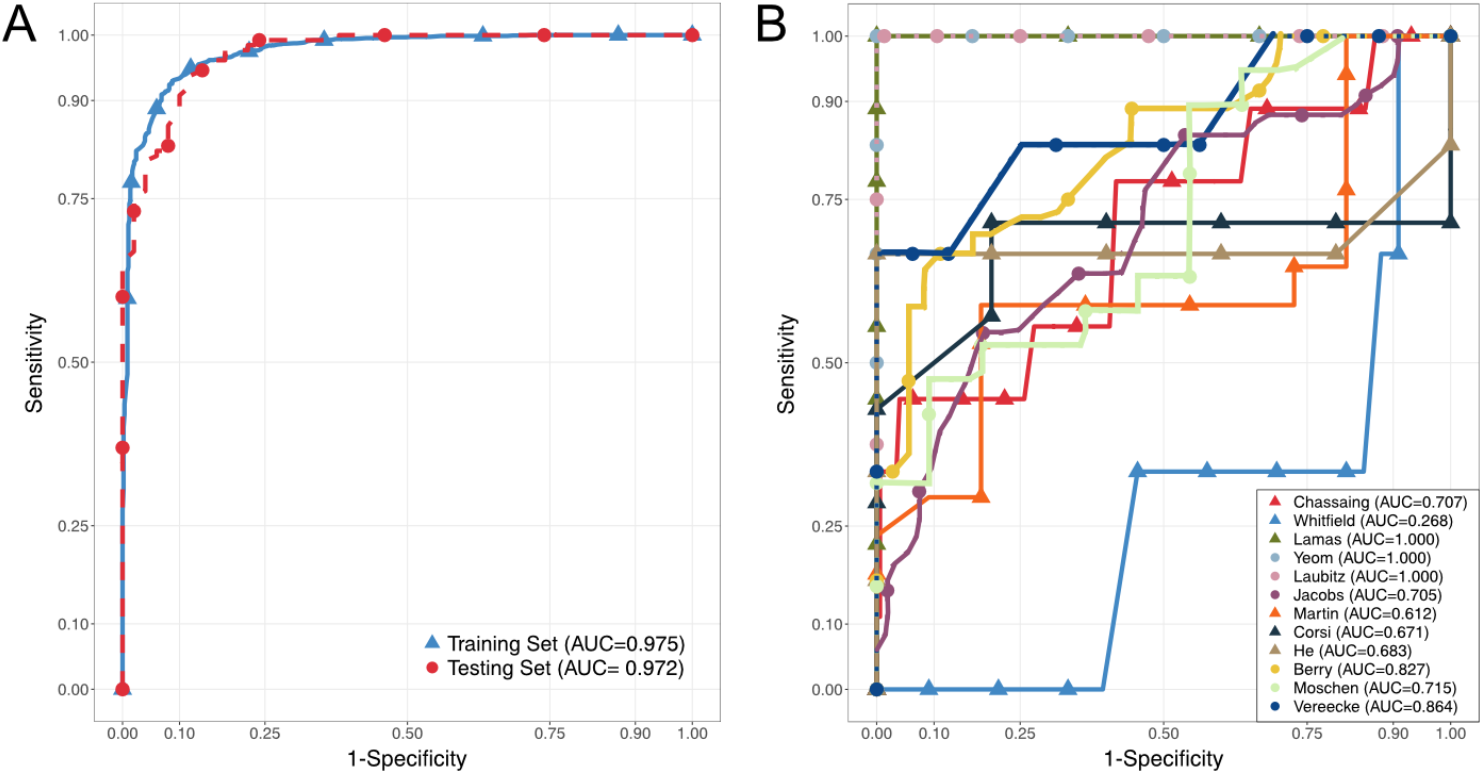
Receiver-Operator Curves demonstrating predictive performance of Random Forest Models, with Area-Under the Curves (AUC) reported. (A) shows cross-validated performance on training set consisting of a random 70% of all samples (blue) and performance on remaining 30% of samples in the testing set (red). (B) shows performance of models trained on (n-1) studies on the study that was left out. Each color-shape and name in the legend refers to the study that was left out of, and predicted by, the model.

To test the generalizability of this approach and whether the individual studies contributed complementary or unique information, we conducted a leave-one-out analysis. One by one, each study was left out, and the samples left out were predicted using a random forest classifier trained on the remaining set of samples (“n-1”). Furthermore, we assessed the cross-validated performance of the “n-1” classifiers, as well as that of models trained on the study left out and tested on the “n-1” set. The performance of the models varied greatly depending on which study was left out, as measured by the AUROC (**Fig 4b**). Samples from Lamas et al. (35), Laubitz et al. (19), and Yeom et al. (22) were perfectly predicted by models trained on all other samples, suggesting that they contained “overlapping” information. All other scenarios had performance worse than the full model, with the Whitfield-Cargile et al. (36) samples having AUROC less than 0.5 (0.268). The worse-than-random prediction of the Whitefield-Cargile et al. samples suggests that it contributes information that runs counter to those in the other studies. Potential reasons for this are considered in the discussion. Assessing from the opposite direction, we also found great variability in the performance of models trained on one study and used to predict the rest. None of these had very high AUROC values, which is likely due to small sample sizes within each study. Indeed, He et al., Corsi et al. and Yeom et al. had few samples, and poorly predicted other studies. On the other hand, Lamas et al. performed well even with small sample sizes. Despite this variability, we found that removing one study did not change the overall model performance, as assessed by repeated cross-validation, with AUROC values staying consistently around 0.9 (**Fig 5**).

**FIG 5:**
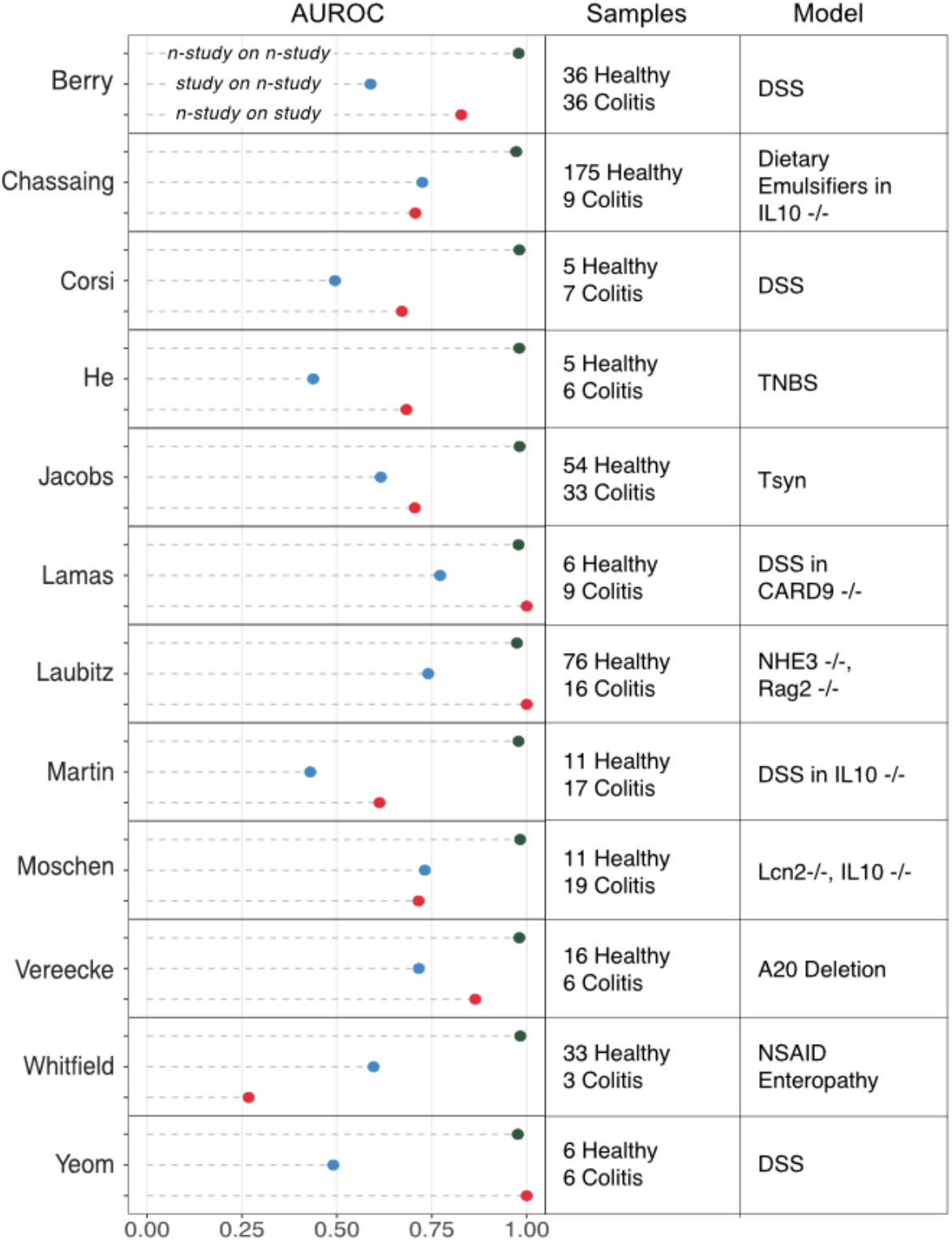
Summary of model performances in terms of Area Under the Receiver-Operator Curve (AUROC) and characteristics for each study. The top line (green) shows cross-validated performance of model when the named study was left out (e.g. for first panel, cross-validated AUROC for model with all studies except Berry et al.). The second line (blue) shows performance of model trained on named study on predicting all other studies (for first panel, AUROC for model trained on Berry et al. and tested on all studies except Berry et al.). The third line (red) shows performance of model trained on all studies except the named one, on predicting the named one (for first panel, AUROC for model trained on all studies except Berry et al. and tested on Berry et al.).

### Variable Selection and “Dysbiosis” Index

As the overall random forest model had a robust predictive performance, we wanted to identify the taxons that contributed to this predictive power. To do this, we used the Boruta feature selection algorithm. This algorithm creates random probes (i.e. taxons with shuffled abundance values across samples), and tests their performance relative to the true features (i.e. taxons with observed abundance values). An iterative process retains only the taxons that are significantly better at prediction than their random counterparts (46). The algorithm yielded 184 taxons that were confirmed as being important for disease status prediction, out of the total 1558 tested (**Supplementary Table 1**). Ninety-eight of these had a higher mean abundance in mice with colitis (colitis-associated taxons), while 86 had a higher mean abundance in healthy mice (health-associated taxons). To narrow this list further in order to identify relevant taxons that were the most prevalent, we only included ones that were present in half or more healthy (if health-associated) or colitis samples (if colitis-associated). This filtering retained 33 taxons, 12 of which were colitis-associated, and 21 of which were health-associated (**Table 2**).

**TABLE 2.**
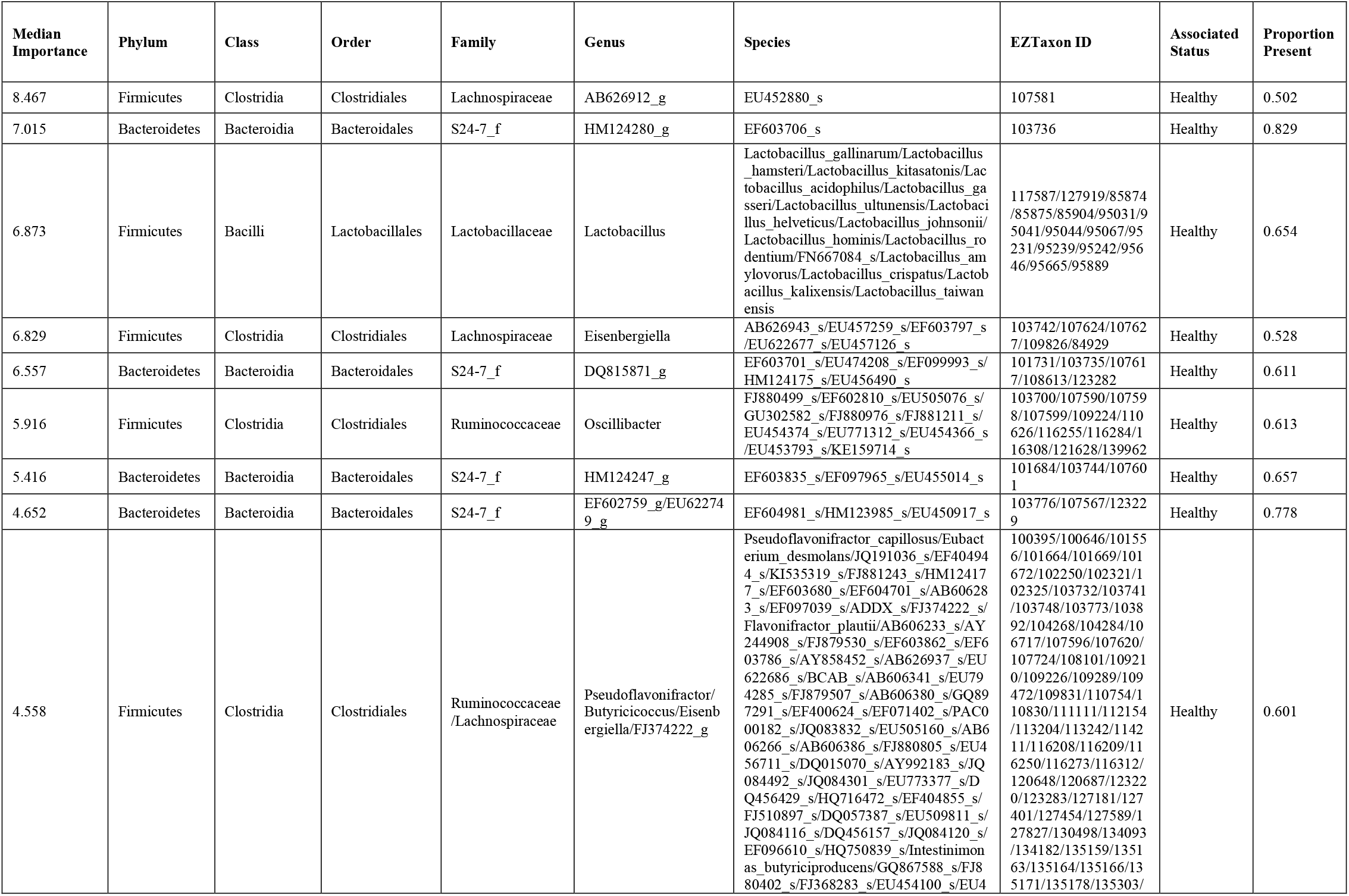
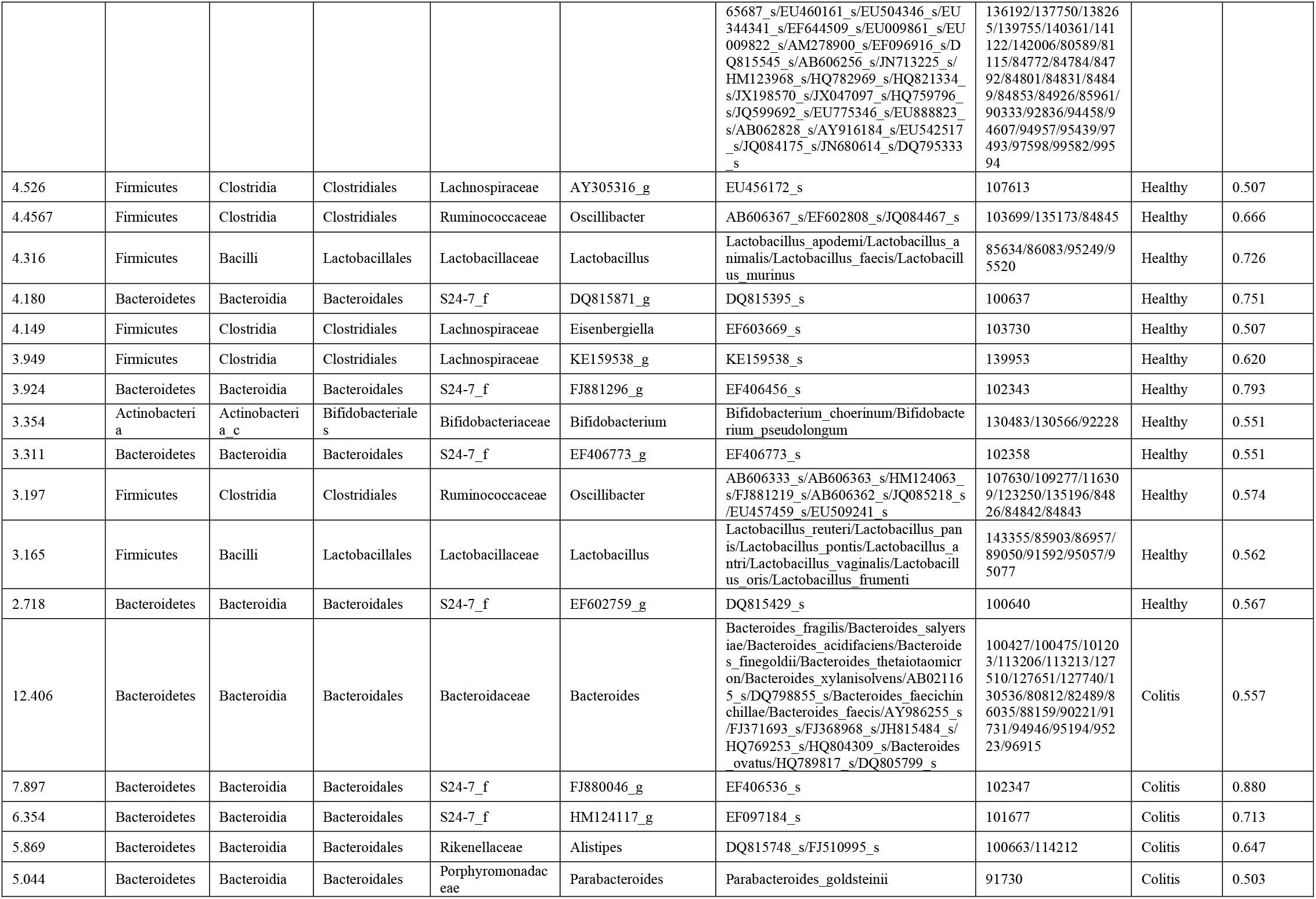
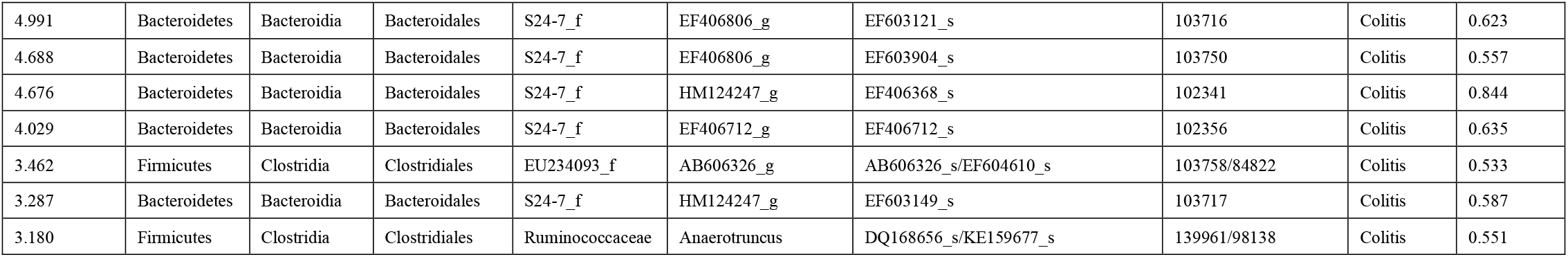
Thirty-three taxa selected through Boruta feature selection for inclusion in dysbiosis index. The median importance is the median importance score calculated by the Boruta algorithm. EZTaxon ID refer to the ID number in the EZTaxon database. Proporotion present refers to the proportion of healthy or colitis samples the taxon was present in.

Most of the microbes (10 of 12) associated with the diseased state were members of the order Bacteroidales, including several species from the abundant but poorly understood S24-7 family. However, microbes associated with the healthy state also included members of the order Bacteroidales (8 of 21) along with a number of Firmicutes (12 of 21 including *Lactobacilli* and *Clostridia sp*.) and one Actinobacteria (*Bifidobacteria*). These findings loosely align with the notion that there may be a bias towards a greater abundance of Firmicutes in the non-diseased state, and that Lactobacilli and Bifidobacteria may be markers of gut health (e.g. both are marketed as probiotics). As it relates to how different Bacteroidales species respond to inflammation and dysbiosis, however, there is clearly much that remains to be understood.

We hypothesized that the combined information contained in the relative abundances of this list of taxons was more relevant for disease status than the Shannon index, BF ratio or abundance of any one taxon. Thus, we created a “dysbiosis index”, which is the log transformed ratio of the relative abundances of colitis-associated taxa to health-associated taxa. When dysbiosis index values was calculated for each sample, we found significantly higher values for colitis samples in 8 studies (t-test, p < 0.05). Two of the remaining had higher, non-significant mean values for mice with colitis, while one had a significantly higher value for healthy samples (**Fig 6**). A random effects model for the pooled data, with study as the random effect, indicated that mice with colitis had significantly higher dysbiosis index values, suggesting that this was a much better indicator of disease status than the other metrics (Shannon diversity index, B/F ratio, proteobacterial abundance) tested above.

**FIG 6:**
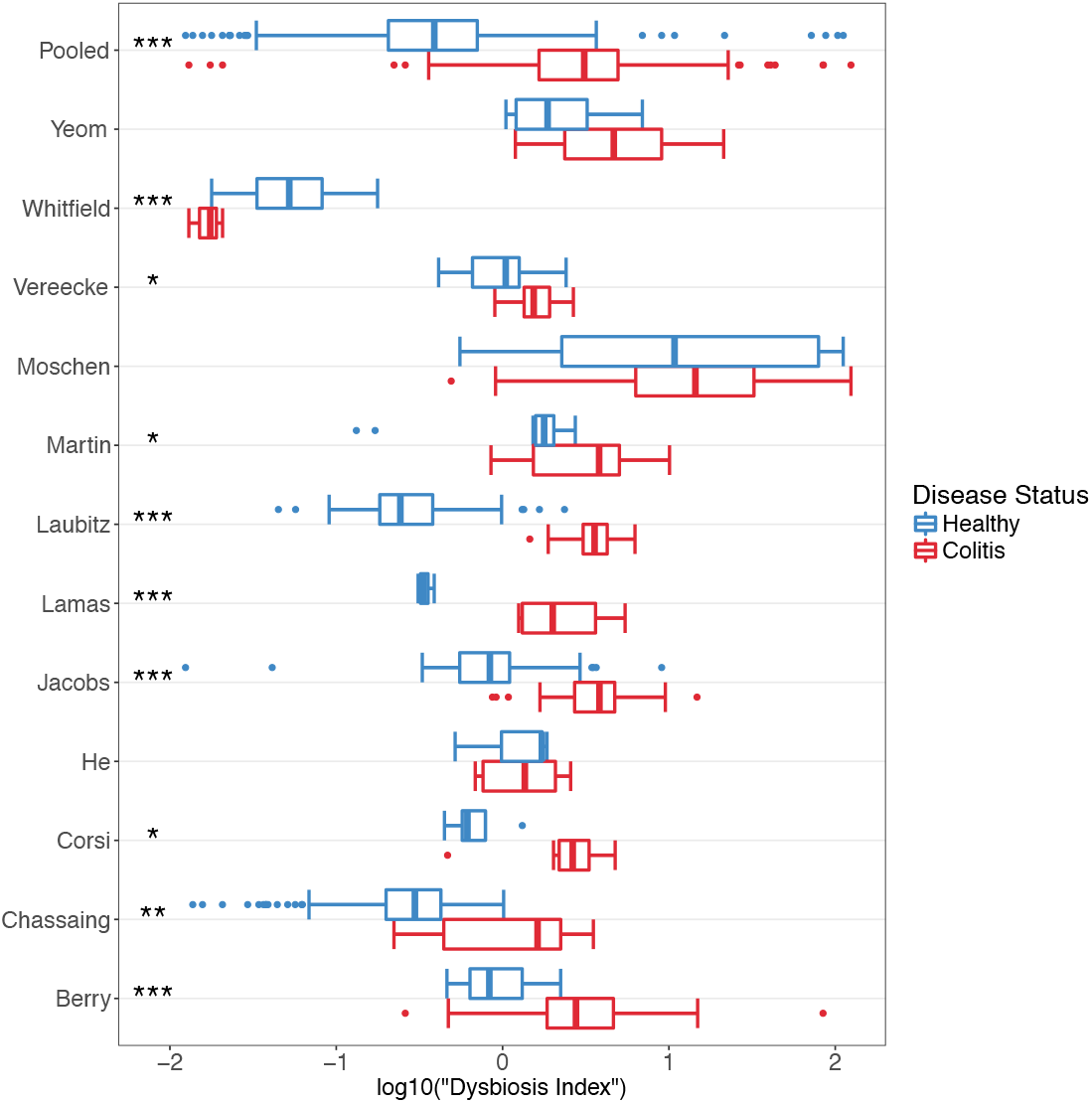
Boxplots showing distribution and statistical significance of “dysbiosis index” within each study and for pooled samples. Blue represents healthy samples and red represents colitic samples. * = p < 0.05, ** = p < 0.01, *** = p < 0.001.

## DISCUSSION

Animal models, especially murine models, have been extensively used in research to understand human health and disease. For IBD in particular, a multitude of chemically induced, cell-transfer, and genetic models in mice have been used to understand aspects of intestinal immunology (18). More recently, there has been increasing interest in how the gut microbiota plays an important role in the onset of IBD in humans and colitis in these murine models. The prevalence of IBD in human populations and its murky etiology, combined with breakthroughs in microbiome research techniques, has fostered a plethora of studies investigating the role of the gut microbiota in this disease. Thus, these studies not only represent a potential for understanding the causes of IBD, but also the role of the microbiome more generally (47). Since it is difficult to interrogate causality and mechanisms of disease onset in humans, murine models provide such work, despite differences in some anatomical features, dietary habits and the microbiota.

In studying complex systems like microbiomes, where there are a nearly endless number of dimensions to be explored in terms of both membership and abundance, summary statistics that enable comparison along one dimension are often attractive. Alpha diversity metrics aim to summarize information regarding the how many members are present, as well as how evenly their abundances are distributed. Here we test the Shannon diversity index and find that even though individual studies can have significant results, they are not consistently in the same direction, and in the pooled data, there is no significant evidence of a relationship. Similarly, the BF ratio and Proteobacteria levels were also found to be inconsistent markers of gut health. Thus, while these one-dimensional metrics have the advantage of simplicity, they should be employed with caution. Moreover, universal generalizations linking a diverse ecosystem to health may be unwarranted in this context.

Another frequently accepted principle is that the functional features of the microbiota can be usefully predicted from 16S level phylogenetic analysis at the family or genus level. Here, again, our study urges caution. While our analysis flagged an increase in specific Bacteroidales during inflammation as a common feature, other closely related Bacteroidales were reduced. Of particular interest are members of the S24-7 family, which are abundant members of the gut microbiota and yet are understudied and poorly understood and may play an important role in the resilience of the microbiota in the face of different abiotic stresses including osmotic-induced diarrhea (48). Given that different members of the S24-7 family were associated with either the disease or healthy state, the assumption that there is functional similarity between microbes at the phylum, family, or even genus level is an oversimplification that may prevent a more nuanced and complete understanding of the specific forces that shape the membership/abundance of gut microbial communities.

When analyzing high-dimensional data, the use of statistical learning techniques can be useful to “pick out” patterns that are not immediately observable – even from highly heterogenous datasets derived from different studies across multiple animal facilities. The results of our leave-one-out analyses indeed highlights the heterogeneity that exists between each of the studies. Given the relatively limited scope of our analysis, we were greatly encouraged by the very high predictive performance of our combined random forest model, which may suggest that there may indeed be some ‘universal’ microbial signatures that enable us to discriminate between healthy and diseased samples across multiple unrelated studies. It would be interesting to investigate what factors contribute to the observed heterogeneity between some studies, especially after having relatively stringent inclusion criteria. This would only be feasible, however, if a much greater number of studies with available data and large samples sizes became available.

We note that one study chosen (Whitfield-Cargile) was a distinct outlier in our analysis from the other eleven studies. Unlike the other genetic and chemical models of dysbiosis included in this meta-analysis, the NSAID-induced inflammation in the Whitfield-Cargile study largely manifests in the distal jejunum and ileum and less the large intestine (i.e. it is not a true “colitis” *per se*). Furthermore, the trigger of dysbiosis used in that study, the NSAID indomethacin, is itself an anti-inflammatory agent that inhibits cyclooxygenase enzymes – which would further change the nature of the resulting inflammation. This indicates that care must be taken when comparing studies that may be superficially similar, but that have important differences in their underlying mechanisms or locations within the host. Accordingly, we believe the fact that the Whitfield-Cargile study performed differently than the other studies, indicates that the dysbiosis index derived from our meta-analysis is robust and able to distinguish colitis from other types of intestinal pathology.

The vast differences in the composition of the human and murine microbiota mean that translation between the two cannot be made directly in terms of individual microbes. On one hand, this means that it might require considerable effort in order to find any potential human counterpart of murine microbes causally implicated in colitis. On the other hand, this has meant that efforts to catalog the human gut microbiome has not helped much with cataloging the murine microbiome and robust human meta-analyses to synthesize evidence on the IBD-microbiome interaction cannot be readily used to guide future studies in mouse models. In this context, the meta-analysis of animal studies can be useful. Generally, such studies are conducted for preclinical animal trials to select robust candidates for clinical trials, and prevent excessive replication (49, 50). However, animal meta-analyses can also play a valuable role in guiding research avenues in murine models of complex systems like microbiomes that are often studied at a macro-level, without a systematic approach to investigating host-microbe or microbe-microbe interactions at a micro level. In our meta-analysis, we find that simple metrics to summarize microbial diversity and composition may not be consistent; we also identify a microbial signature of disease that is relatively robust across studies, and report a list of microbes that may be good candidates for focused isolation and characterization efforts.

Most importantly, through the quantitative synthesis of published literature, we identified a number of organisms that seem to be consistently associated with health or disease in murine models of IBD. The colitis-associated microbes are likely to be good candidates for screening in mono-association or infection studies, whereas the healthy-associated ones are likely to be good candidates for probiotic studies. Many of the identified strains are phylotypes that are yet to be isolated. In a rapidly evolving field where mice microbiomes have been under increasing attention for systematic approaches to cataloguing and strain isolation, this study provides a tool that can be used to prioritize efforts.

Finally, our experience suggests that trying to standardize microbiome studies and make the data publicly accessible is of paramount importance. One of the main time-consuming steps in our analysis was the custom processing of datasets generated by a diversity of sequencing approaches, and the resolution of our taxonomic classification was limited by the diversity of 16S primers used. While we identified 44 studies of interest for the meta-analysis, we were only able to obtain data from 12, suggesting that only a small fraction of published sequencing data is actually deposited in a publicly accessible system. Journal policies on data-sharing can help rectify this.

## METHODS

### Study Search and Inclusion

To identify studies that investigated the murine intestinal microbiota in the context of intestinal inflammation, we conducted a systematic search of NCBI PubMed for articles that contained terms relating to microbiota, intestinal inflammation and murine models in the title and abstract, published between 2012 and 2016, and was not a review. We followed the preferred reporting items for systematic reviews and meta-analyses (PRISMA) guidelines to limit inclusion bias (51). The detailed search term was: “((microbiota[Title/Abstract] OR microbiome[Title/Abstract]) AND (colitis[Title/Abstract] OR (inflammation[Title/Abstract] AND (mucosa*[Title/Abstract] OR epitheli*[Title/Abstract] OR colon*[Title/Abstract] OR gut[Title/Abstract] OR intestin*[Title/Abstract])))) AND (“2012/01/01”[PDAT]: “2016/12/30”[PDAT]) AND (mice[Title/Abstract] OR mouse[Title/Abstract] OR murine[Title/Abstract]) NOT review[Publication Type]”. We screened the title and abstracts of 816 articles yielded by the search for eligibility. In addition, we obtained three studies from knowledge of published literature, and a currently unpublished dataset of an IL-10 knockout model of colitis from a collaborator.

Seventy-nine full-text articles were then assessed, and 44 were retained to be checked for data availability. Ten studies had read files accessible in Sequence Read Archive, European Nucleotide Archive, MG-RAST or personal collaboration; 32 studies had no publicly available data or metadata, and only two provided access to data by the time of publication after contact. Studies were excluded if they: did not do a 16S sequence-based analysis of the microbiome; used a synthetic microbiota (e.g. Altered Schaedler Flora, humanized); only had outcomes of low-grade inflammation or aging associated “inflammaging”; used pathogenic infection for inflammation; used bacterial treatment (e.g. probiotics); used antibiotic treatment; used non-murine models; did not have an outcome of colonic inflammation; did not have non-colitis controls; was not a primary article; or that failed to assess the microbiota before onset of inflammation. We had the final exclusion criteria because many studies aim to answer the question of whether a particular microbiota is associated with increased susceptibility to colitis; however, we were interested in a microbial signature associated with the *onset* of colitis. We retained articles if they contained a subset of samples that were eligible, contingent on non-colitis controls being present. Within the included studies, we excluded any samples that met relevant exclusion criteria outlined above (e.g. low-grade inflammation, antibiotic treatment).

### Data Processing

For each study, we used a standardized bioinformatics pipeline to generate counts for taxonomic groups. Quality filtering criteria was determined on a study-by-study basis depending on the sequencing platform used and inspection of read quality (52, 53). Reads were filtered and resolved to amplicon sequence variants (ASVs) using the DADA2 pipeline. The advantage of using DADA2 over traditional clustering methods is that it resolves differences of as little as one nucleotide to determine exact sequences based on an error model for the sequencing run (54). Resolved paired-end sequences were merged where applicable, chimeras removed and a RSV-abundance tables built (equivalent to OTU-tables).

RSVs were assigned microbial taxonomies using a custom R script (55). Sequences were searched against the EZTaxon database (January 2017 Update) (30) using vsearch (56) and assigned the taxonomy (up to species level) of the highest identity match >97%. Tied hits were assigned ambiguous classification (e.g. Shigella/Escherichia). Use of EZTaxon allows classification of uncultured bacteria because it has unique identifiers assigned to manually curated phylotypes. RSVs classified as “Streptophyta” at phylum level were filtered out. RSV counts were standardized by calculating relative abundances. Within each study, RSVs that did not make up more than 0.2% of the community in at least one sample, as well as taxonomic groups that were detectable in only one sample, were removed.

To enable merging of datasets from different studies, that used different 16S regions, RSVs unclassified at the species level were removed. Next, RSV abundance tables were collapsed to the highest taxonomic resolution possible (e.g. x/y from one study was collapsed into x/y/z group if another study could not resolve between x/y/z). Data handling, merging and filtering was done using the phyloseq package (57).

### Alpha and Beta Diversity Analysis

For each sample, we estimated the Shannon diversity index using the vegan package (58), as well as the ratio of Bacteroidetes abundance to Firmicutes abundance (BF ratio). Following normalizing transformations, t-tests were used to detect significant differences between healthy and mice with colitis, within each study. For the pooled data, a linear random effects model (implemented using the lme4 package (59)) with random slopes and intercepts was used to determine if there was a statistically significant association between the measures and disease status.

Diversity across samples was investigated using Principal Coordinates Analysis (PCoA) and permutational analysis of variance (PERMANOVA) (adonis function; 999 permutations) using the vegan package. This was done separately for each study, as well as the pooled data.

### Random Forest Models

A random forest (RF) model was built with the pooled data using the caret package (60) (10-fold cross validated, repeated 5 times; 250 trees), and area under the receiver operating characteristic curve (AUROC) was sued to assess model performance in discriminating healthy from diseased mice. To investigate the generalizability of using RF models, we conducted a leave-one-out analysis, repeatedly building models on data from n-1 studies, and testing model performance in predicting the study left out, using AUROC.

### Variable Selection and “Dysbiosis” Index

To identify which taxonomic groups are most important in discriminating between healthy and diseased mice within a random forest framework, the Boruta feature selection algorithm was used with 500 runs using the Boruta package (46). Next, the selected taxa were labelled as being associated with healthy or diseased status based on whether they had higher mean abundance in healthy or diseased mice, respectively (statistical significance not considered). This labelled list was pruned to contain only taxa present in 50% of diseased or healthy mice. The dysbiosis index was calculated by log transforming the ratio of the colitis-associated taxa to the health-associated taxa.

As with the alpha diversity metrics, within each study, t-tests were used to determine the utility of the index in discriminating between healthy and diseased mice, and a similar random effects model was used for the pooled data.

All data was visualized using the ggplot2 package (61).

## ACKNOWLEDGEMENTS

We want to thank all the authors who ensured that their data was publicly available or shared the data upon request. This work was funded by the Canadian Institutes of Health Research (CIHR grant 144628).

We also wish to acknowledge this land on which the University of Toronto operates. For thousands of years it has been the traditional land of the Huron-Wendat, the Seneca, and most recently, the Mississaugas of the Credit River. Today, this meeting place is still the home to many Indigenous people from across Turtle Island and we are grateful to have the opportunity to work on this land.

## REFERENCES

1. Marchesi JR, Ravel J. 2015. The vocabulary of microbiome research: a proposal. Microbiome 3:31.

2. Flint HJ, Scott KP, Duncan SH, Louis P, Forano E. 2012. Microbial degradation of complex carbohydrates in the gut. Gut Microbes 3:289–306.

3. Tremaroli V, Bäckhed F. 2012. Functional interactions between the gut microbiota and host metabolism. Nature 489:242–249.

4. Round JL, Mazmanian SK. 2009. The gut microbiota shapes intestinal immune responses during health and disease. Nat Rev Immunol 9:313–323.

5. Petersen C, Round JL. 2014. Defining dysbiosis and its influence on host immunity and disease. Cell Microbiol 16:1024–1033.

6. Kostic AD, Xavier RJ, Gevers D. 2014. The Microbiome in Inflammatory Bowel Disease: Current Status and the Future Ahead. Gastroenterology 146:1489–1499.

7. Shanahan F. 2002. Crohn’s disease. Lancet Lond Engl 359:62–69.

8. Farrell RJ, Peppercorn MA. 2002. Ulcerative colitis. Lancet Lond Engl 359:331–340.

9. Jovel J, Patterson J, Wang W, Hotte N, O’Keefe S, Mitchel T, Perry T, Kao D, Mason AL, Madsen KL, others. 2016. Characterization of the Gut Microbiome Using 16S or Shotgun Metagenomics. Front Microbiol 7.

10. Hiergeist A, Reischl U, Gessner A. 2016. Multicenter quality assessment of 16S ribosomal DNA-sequencing for microbiome analyses reveals high inter-center variability. Int J Med Microbiol 306:334–342.

11. Walters WA, Xu Z, Knight R. 2014. Meta-analyses of human gut microbes associated with obesity and IBD. FEBS Lett 588:4223–4233.

12. Shah MS, DeSantis TZ, Weinmaier T, McMurdie PJ, Cope JL, Altrichter A, Yamal J-M, Hollister EB. 2017. Leveraging sequence-based faecal microbial community survey data to identify a composite biomarker for colorectal cancer. Gut gutjnl–2016.

13. Sze MA, Schloss PD. 2016. Looking for a signal in the noise: revisiting obesity and the microbiome. MBio 7:e01018–16.

14. Hugenholtz F, de Vos WM. 2018. Mouse models for human intestinal microbiota research: a critical evaluation. Cell Mol Life Sci 75:149–160.

15. Hörmannsperger G, Schaubeck M, Haller D. 2015. Intestinal Microbiota in Animal Models of Inflammatory Diseases. ILAR J 56:179–191.

16. Hildebrand F, Nguyen TLA, Brinkman B, Yunta RG, Cauwe B, Vandenabeele P, Liston A, Raes J. 2013. Inflammation-associated enterotypes, host genotype, cage and inter-individual effects drive gut microbiota variation in common laboratory mice. Genome Biol 14:R4.

17. Kiesler P, Fuss IJ, Strober W. 2015. Experimental models of inflammatory bowel diseases. Cell Mol Gastroenterol Hepatol 1:154–170.

18. Mizoguchi A. 2012. Animal models of inflammatory bowel disease, p. 263–320. In Progress in molecular biology and translational science. Elsevier.

19. Laubitz D, Harrison CA, Midura-Kiela MT, Ramalingam R, Larmonier CB, Chase JH, Caporaso JG, Besselsen DG, Ghishan FK, Kiela PR. 2016. Reduced epithelial Na+/H+ exchange drives gut microbial dysbiosis and promotes inflammatory response in T cell-mediated murine colitis. PloS One 11:e0152044.

20. Moschen AR, Gerner RR, Wang J, Klepsch V, Adolph TE, Reider SJ, Hackl H, Pfister A, Schilling J, Moser PL, Kempster SL, Swidsinski A, Orth Höller D, Weiss G, Baines JF, Kaser A, Tilg H. 2016. Lipocalin 2 Protects from Inflammation and Tumorigenesis Associated with Gut Microbiota Alterations. Cell Host Microbe 19:455–469.

21. Berry D, Schwab C, Milinovich G, Reichert J, Mahfoudh KB, Decker T, Engel M, Hai B, Hainzl E, Heider S, others. 2012. Phylotype-level 16S rRNA analysis reveals new bacterial indicators of health state in acute murine colitis. ISME J 6:2091–2106.

22. Yeom Y, Kim B-S, Kim S-J, Kim Y. 2016. Sasa quelpaertensis leaf extract regulates microbial dysbiosis by modulating the composition and diversity of the microbiota in dextran sulfate sodium-induced colitis mice. BMC Complement Altern Med 16:303.

23. Dziarski R, Park SY, Kashyap DR, Dowd SE, Gupta D. 2016. Pglyrp-Regulated Gut Microflora Prevotella falsenii, Parabacteroides distasonis and Bacteroides eggerthii Enhance and Alistipes finegoldii Attenuates Colitis in Mice. PloS One 11:e0146162.

24. Sokol H, Pigneur B, Watterlot L, Lakhdari O, Bermúdez-Humarán LG, Gratadoux J-J, Blugeon S, Bridonneau C, Furet J-P, Corthier G. 2008. Faecalibacterium prausnitzii is an anti-inflammatory commensal bacterium identified by gut microbiota analysis of Crohn disease patients. Proc Natl Acad Sci 105:16731–16736.

25. Nell S, Suerbaum S, Josenhans C. 2010. The impact of the microbiota on the pathogenesis of IBD: lessons from mouse infection models. Nat Rev Microbiol 8:564.

26. Gkouskou K, Deligianni C, Tsatsanis C, Eliopoulos AG. 2014. The gut microbiota in mouse models of inflammatory bowel disease. Front Cell Infect Microbiol 4.

27. Xiao L, Feng Q, Liang S, Sonne SB, Xia Z, Qiu X, Li X, Long H, Zhang J, Zhang D. 2015. A catalog of the mouse gut metagenome. Nat Biotechnol 33:1103.

28. Lagkouvardos I, Pukall R, Abt B, Foesel BU, Meier-Kolthoff JP, Kumar N, Bresciani A, Martínez I, Just S, Ziegler C. 2016. The Mouse Intestinal Bacterial Collection (miBC) provides host-specific insight into cultured diversity and functional potential of the gut microbiota. Nat Microbiol 1:16131.

29. Callahan BJ, McMurdie PJ, Holmes SP. 2017. Exact sequence variants should replace operational taxonomic units in marker-gene data analysis. ISME J 11:2639–2643.

30. Kim O-S, Cho Y-J, Lee K, Yoon S-H, Kim M, Na H, Park S-C, Jeon YS, Lee J-H, Yi H, Won S, Chun J. 2012. Introducing EzTaxon-e: a prokaryotic 16S rRNA gene sequence database with phylotypes that represent uncultured species. Int J Syst Evol Microbiol 62:716–721.

31. Shin N-R, Whon TW, Bae J-W. 2015. Proteobacteria: microbial signature of dysbiosis in gut microbiota. Trends Biotechnol 33:496–503.

32. Moher D, Liberati A, Tetzlaff J, Altman DG, Group TP. 2009. Preferred Reporting Items for Systematic Reviews and Meta-Analyses: The PRISMA Statement. PLOS Med 6:e1000097.

33. Jacobs JP, Lin L, Goudarzi M, Ruegger P, McGovern DP, Fornace Jr AJ, Borneman J, Xia L, Braun J. 2016. Microbial, metabolomic, and immunologic dynamics in a relapsing genetic mouse model of colitis induced by T-synthase deficiency. Gut Microbes 00–00.

34. Chassaing B, Koren O, Goodrich JK, Poole AC, Srinivasan S, Ley RE, Gewirtz AT. 2015. Dietary emulsifiers impact the mouse gut microbiota promoting colitis and metabolic syndrome. Nature 519:92.

35. Lamas B, Richard ML, Leducq V, Pham H-P, Michel M-L, Da Costa G, Bridonneau C, Jegou S, Hoffmann TW, Natividad JM, others. 2016. CARD9 impacts colitis by altering gut microbiota metabolism of tryptophan into aryl hydrocarbon receptor ligands. Nat Med 22:598–605.

36. Whitfield-Cargile CM, Cohen ND, Chapkin RS, Weeks BR, Davidson LA, Goldsby JS, Hunt CL, Steinmeyer SH, Menon R, Suchodolski JS, others. 2016. The microbiota-derived metabolite indole decreases mucosal inflammation and injury in a murine model of NSAID enteropathy. Gut Microbes 1–16.

37. Sassone-Corsi M, Nuccio S-P, Liu H, Hernandez D, Vu CT, Takahashi AA, Edwards RA, Raffatellu M. 2016. Microcins mediate competition among Enterobacteriaceae in the inflamed gut. Nature 540:280–283.

38. Vereecke L, Vieira-Silva S, Billiet T, van Es JH, Mc Guire C, Slowicka K, Sze M, van den Born M, De Hertogh G, Clevers H, Raes J, Rutgeerts P, Vermeire S, Beyaert R, van Loo G. 2014. A20 controls intestinal homeostasis through cell-specific activities. Nat Commun 5:5103.

39. He Q, Li X, Liu C, Su L, Xia Z, Li X, Li Y, Li L, Yan T, Feng Q, others. 2016. Dysbiosis of the fecal microbiota in the TNBS-induced Crohn’s disease mouse model. Appl Microbiol Biotechnol 100:4485–4494.

40. Goodrich JK, Di Rienzi SC, Poole AC, Koren O, Walters WA, Caporaso JG, Knight R, Ley RE. 2014. Conducting a microbiome study. Cell 158:250–262.

41. Bray JR, Curtis JT. 1957. An ordination of the upland forest communities of southern Wisconsin. Ecol Monogr 27:325–349.

42. Anderson MJ. 2005. PERMANOVA: a FORTRAN computer program for permutational multivariate analysis of variance. Dep Stat Univ Auckl N Z 24.

43. Duvallet C, Gibbons SM, Gurry T, Irizarry RA, Alm EJ. 2017. Meta-analysis of gut microbiome studies identifies disease-specific and shared responses. Nat Commun 8:1784.

44. Vázquez-Baeza Y, Hyde ER, Suchodolski JS, Knight R. 2016. Dog and human inflammatory bowel disease rely on overlapping yet distinct dysbiosis networks. Nat Microbiol 1:16177.

45. Breiman L. 2001. Random forests. Mach Learn 45:5–32.

46. Kursa MB, Rudnicki WR. 2010. Feature selection with the Boruta package. J Stat Softw 36:1–13.

47. Huttenhower C, Kostic AD, Xavier RJ. 2014. Inflammatory Bowel Disease as a Model for Translating the Microbiome. Immunity 40:843–854.

48. Tropini C, Moss EL, Merrill BD, Ng KM, Higginbottom SK, Casavant EP, Gonzalez CG, Fremin B, Bouley DM, Elias JE, Bhatt AS, Huang KC, Sonnenburg JL. 2018. Transient Osmotic Perturbation Causes Long-Term Alteration to the Gut Microbiota. Cell 173:1742–1754.e17.

49. Vesterinen HM, Sena ES, Egan KJ, Hirst TC, Churolov L, Currie GL, Antonic A, Howells DW, Macleod MR. 2014. Meta-analysis of data from animal studies: a practical guide. J Neurosci Methods 221:92–102.

50. Hooijmans CR, IntHout J, Ritskes-Hoitinga M, Rovers MM. 2014. Meta-analyses of animal studies: an introduction of a valuable instrument to further improve healthcare. ILAR J 55:418–426.

51. Liberati A, Altman DG, Tetzlaff J, Mulrow C, Gøtzsche PC, Ioannidis JPA, Clarke M, Devereaux PJ, Kleijnen J, Moher D. 2009. The PRISMA Statement for Reporting Systematic Reviews and Meta-Analyses of Studies That Evaluate Health Care Interventions: Explanation and Elaboration. PLOS Med 6:e1000100.

52. Callahan BJ, Sankaran K, Fukuyama JA, McMurdie PJ, Holmes SP. 2016. Bioconductor workflow for microbiome data analysis: from raw reads to community analyses. F1000Research 5:1492.

53. Callahan B. DADA2 Pipeline Tutorial (1.8).

54. Callahan BJ, McMurdie PJ, Rosen MJ, Han AW, Johnson AJA, Holmes SP. 2016. DADA2: High-resolution sample inference from Illumina amplicon data. Nat Methods 13:581–583.

55. Team RC. 2014. R: A language and environment for statistical computing. R Foundation for Statistical Computing, Vienna, Austria. 2013.

56. Rognes T, Flouri T, Nichols B, Quince C, Mahé F. 2016. VSEARCH: a versatile open source tool for metagenomics. PeerJ 4:e2584.

57. McMurdie PJ, Holmes S. 2013. phyloseq: An R Package for Reproducible Interactive Analysis and Graphics of Microbiome Census Data. PLOS ONE 8:e61217.

58. Oksanen J, Kindt R, Legendre P, O’Hara B, Stevens MHH, Oksanen MJ, Suggests M. 2007. The vegan package. Community Ecol Package 10:631–637.

59. Bates D, Mächler M, Bolker B, Walker S. 2014. Fitting linear mixed-effects models using lme4. ArXiv Prepr ArXiv14065823.

60. Kuhn M. 2008. Caret package. J Stat Softw 28:1–26.

61. Wickham H. 2010. ggplot2: elegant graphics for data analysis. J Stat Softw 35:65–88.

