## Supplementary figures and images for "Meta-analysis identifies microbial signatures of disease in murine models of inflammatory bowel disease"

### Supplemental Figure 1

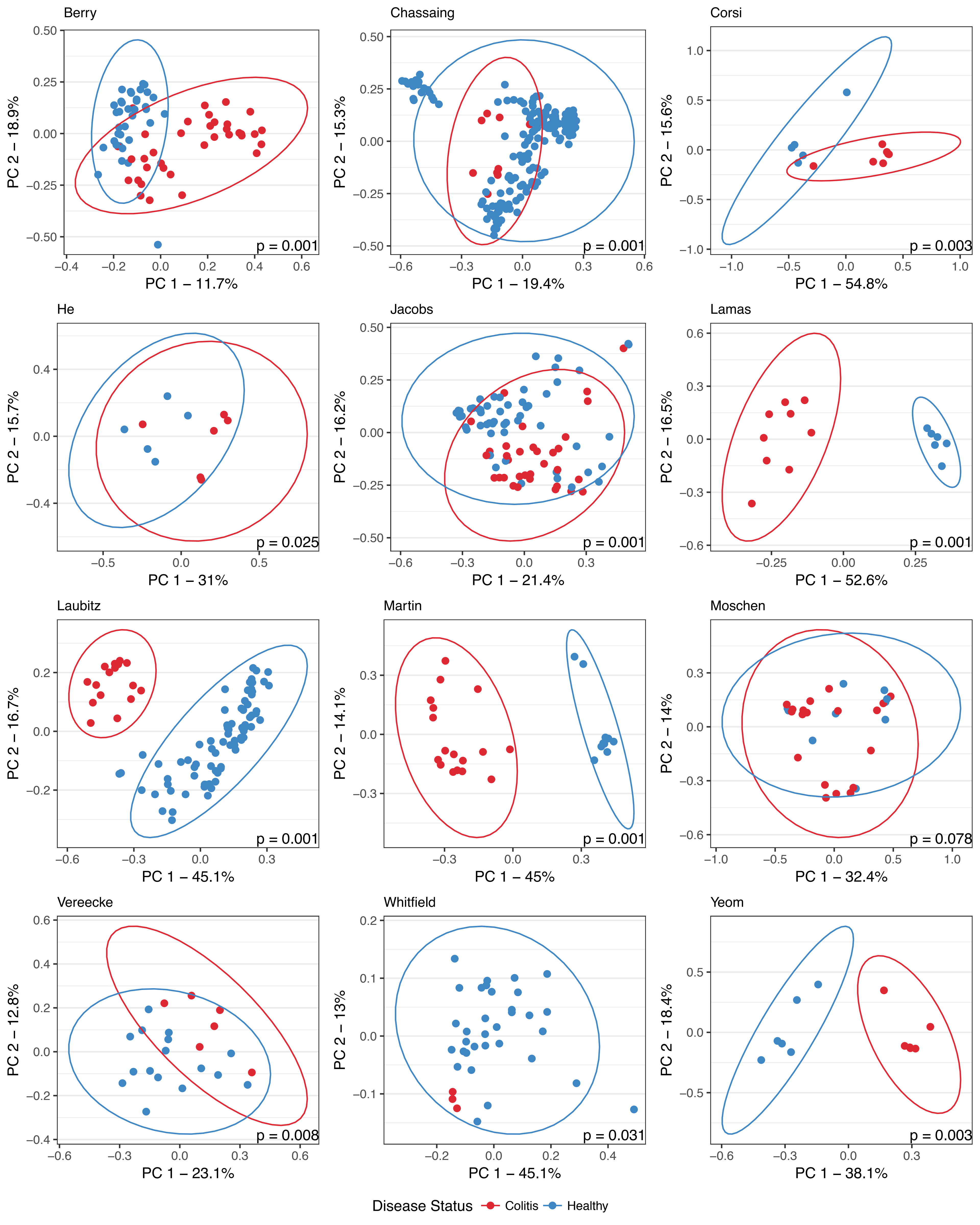
